# Evaluating the Anti-Tumor Efficacy and Safety of AEV01 in Glioblastoma and Hepatocellular Carcinoma: An *In Vitro* and *In Vivo* Approach

**DOI:** 10.1101/2024.07.22.604571

**Authors:** Indrani Biswas, Daisy Precilla S, Aravinda Kumar, MM Sekhar, Renu Medasani, Anitha TS

**Affiliations:** Mahatma Gandhi Medical Advanced Research Institute, Sri Balaji Vidyapeeth University, Puducherry; Department of Pharmacology, Pondicherry Institute of Medical Sciences, Ganapathichettikulam, Puducherry; Astrel Genome Ltd, Hyderabad, India; Department of Biochemistry & Molecular Biology, School of Life Sciences, Pondicherry University, Puducherry

**Keywords:** AEV01, Glioblastoma, Hepatocellular Carcinoma, Inflammation, *in vitro*, *in vivo*

## Abstract

Gliomas and hepatocellular carcinoma (HCC) are aggressive cancers with poor prognoses, often leading to less than a year of survival. Therapeutic resistance underscores the need for novel therapeutic strategies. AEV01, derived from *Picrorhiza kurroa*, has shown promise as a potential anticancer agent. In this context, the current study aimed to evaluate the anti-tumor efficacy and safety profile of AEV01 both *in vitro* and *in vivo* in glioblastoma and HCC models. Briefly, cytotoxicity and apoptosis were assessed using MTT assays and AO/EtBr staining, while ELISA and immunofluorescence measured inflammatory markers, TP53 and CD36 expression. *In vivo*, ELISA was performed against the inflammatory and tumor suppressor markers while, histopathological analysis assessed tumor morphology and organ toxicity. AEV01 exhibited dose-dependent cytotoxicity against U-87 MG glioblastoma and HepG2 liver cancer cells, with optimal concentrations at 400 µg and 300 µg respectively. Treatment downregulated inflammatory markers, CD36 expression and concomitantly increased TP53 expression. Xenograft models depicted similar results, with reduced tumor markers expression, reduced tissue architecture, and no significant organ toxicity. Thus, AEV01 demonstrated potent anti-tumor activity with a favorable safety profile, suggesting its potential as a novel therapeutic agent for gliomas and HCC, warranting further clinical investigation.

## Introduction

Gliomas and hepatocellular carcinoma (HCC) represent alarming challenges in the field of oncology, characterized by their aggressive nature, poor prognosis, and limited response to existing therapies ^1,2^. Gliomas, particularly glioblastoma multiforme (GBM), have a dismal median survival time of approximately 12-15 months despite the use of surgery, radiation, and chemotherapy ^3^. Similarly, HCC, the most common form of primary liver cancer, is associated with high recurrence rates and a median survival of less than a year in advanced stages, even with the application of current standard-of-care treatments such as sorafenib and Lenvatinib ^4^. The limited efficacy of these therapeutic options, combined with the rapid development of resistance and the scarcity of successful clinical trials, underscores the critical need for novel therapeutic strategies.

Orthotopic cell-derived tumor xenograft (CDX) models have provided valuable insights into the complex biology of glioblastoma (GB) and HCC, aiding in the identification of new therapeutic targets ^5^. These models, which more accurately mimic the human tumor microenvironment compared to subcutaneous models, have been instrumental in preclinical research aimed at identifying and validating new treatment approaches ^6^.

In response to the urgent need for innovative cancer therapies, AEV01, a novel drug product developed by Astrel Genome Ltd, Hyderabad, India, emerges as a promising candidate. AEV01 is a specialized extract derived from the roots and rhizomes of *Picrorhiza kurroa*, an alpine herb native to the Himalayan region. *Picrorhiza kurroa*, known as “Kutki” in traditional Indian medicine, has a long-standing reputation for its therapeutic properties, including anti-inflammatory, antioxidant, hepatoprotective, immunostimulant, neuroprotective, and anticancer effects ^7,8^. These properties are attributed to the herb’s bioactive compounds, such as picroside I and II, which have been shown to modulate various biological pathways involved in inflammation, oxidative stress, and immune responses ^9,10^.

Recent scientific evidence highlights the potential of AEV01 in cancer treatment. In a case study involving a 68-year-old patient with metastatic pancreatic cancer, AEV01 administration led to significant clinical improvements, including pain relief, increased energy levels, and enhanced recovery from chemotherapy. Remarkably, the patient exhibited normalized blood markers, weight gain, and fully rejuvenated T-cell function over a two-month period, culminating in a return to normal, active life (unpublished data). Such outcomes, though preliminary, suggest that AEV01 may exert profound effects on immune modulation and tumor biology.

Furthermore, a randomized, placebo-controlled, double-blinded study conducted by Balan et al. demonstrated the safety and efficacy of AEV01 in enhancing immune responses in COVID-19 patients ^7^. The study revealed that AEV01 significantly reduced the neutrophil-to-lymphocyte ratio (NLR) and modulated CD4+/CD8+ T-cell populations, leading to improved vaccination efficacy in aged adults ^7^. These findings underscore the potential of AEV01 not only in viral infections but also in broader therapeutic applications, including cancer treatment, where immune modulation plays a critical role.

Given these promising preliminary results, this study aims to systematically investigate the anti-tumor properties of AEV01 in U-87 MG (Human GB) and HepG2/Hep3B (Human Liver Cancer) cells. The research involves both *in vitro* experiments to assess the cytotoxic and anti-proliferative effects of AEV01 on cancer cells and *in vivo* studies using xenograft-induced animal models to evaluate its efficacy in reducing tumor growth and improving survival outcomes. By integrating preclinical evidence with emerging clinical observations, this study seeks to validate AEV01 as a potential therapeutic agent in the treatment of gliomas and HCC, addressing a critical need in oncology.

## Methods

### • Chemicals

MTT (3-(4,5-dimethylthiazolyl-2)-2,5-diphenylte-trazolium bromide), DAPI (4′,6-diamidino-2-phenylindole), and all the other reagents used for the experiments were obtained from Sigma (USA) and HiMedia (Mumbai, India).

### • Cell Line and culture conditions

HEK-293T Human embryonic kidney cell line (RRID: CVCL_0063) (Passage No: 10), U-87 MG human GB cell line (RRID: CVCL_5J15) (Passage Number: 58), HepG2 human liver cancer cell line (RRID: CVCL_0027) (Passage Number: 35), Hep3B human liver cancer cell line (RRID: CVCL_0326) (Passage Number: 24) and WRL-68 liver senescence cells (RRID: CVCL_0581) (Passage Number: 68) were procured from National Centre for Cell Sciences (NCCS) (Pune, India). HEK-293T, HepG2, Hep3B and WRL-68 cells were grown in Dulbecco’s Modified Eagle’s Medium (DMEM) with 10% FBS and 1% antibiotic solution (100 U/ml penicillin and 100 μg/ml streptomycin), while, U-87 MG human GB cell lines were maintained in Minimal Essential Medium Eagle (EMEM) with 10% FBS and 1% antibiotic solution (100 U/ml penicillin and 100 μg/ml streptomycin). The cells were maintained at 37°C under humidified 5% CO_2_ - 95% O_2_. At their logarithmic growth phase, the cells were trypsinized, counted using trypan blue dye-exclusion method and resuspended in 1X-PBS to a final concentration of 1×10^6^ cells per 3 µl of PBS.

### • Preparation of AEV01 Drug for Experiments

The main stock solution of the AEV01 drug was supplied at a concentration of 100 mg/ml dissolved in the vehicle. Working stock solutions were then prepared at a concentration of 1 mg/ml and employed for the experiments. These aliquots were stored at −20°C. On the day of the experiments, the AEV01 drug was dissolved in the culture medium for application.

### • MTT Assay for Cell Viability

To determine the cytotoxic effect of AEV01 on cancer and non-cancerous cells, MTT was performed on U-87 MG, HepG2 and HEK293T cells. Briefly, during their logarithmic growth phase, the cells were harvested using 0.25% Trypsin and 0.53 mM EDTA, then counted using the trypan-blue dye exclusion assay. Cells were seeded in 96-well plates at a density of 5×10³ cells per well. After 24 hours, the cells were washed with 1X-PBS and treated with varying concentrations of the drug and vehicle control (VC) ranging from 100 to 700 µg for an additional 24 hours. Following treatment, the media was aspirated, and 20 µl of MTT reagent (5 mg/ml) was added to each well. The cells were then incubated in the dark at 37°C for 4 hours. The resulting purple formazan crystals were dissolved by adding DMSO to each well. Absorbance at 570 nm was measured using an ELISA-Multiplate reader (Molecular Devices Spectra-Max M5, USA). The absorbance values were normalized to the control group for relative quantification. The percentage viability was calculated using the formula:

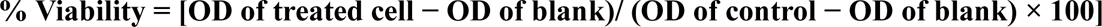

### • AO/EtBr Dual Staining for Apoptosis Assessment

Acridine orange/ethidium bromide (AO/EtBr) dual staining was performed to assess morphological changes induced by AEV01 in U-87 MG and HepG2 cancer cells. Cells were seeded at a density of 20×10³ cells per well in a 24-well plate and treated with the drug for 24 hours at 37°C. Following incubation, the cells were washed with 1X PBS and stained with 200 µl of AO/EtBr solution (100 μg/ml each) at room temperature for 5 minutes. The stained cells were then visualized using a fluorescent microscope (Axiovert 40 CFL, Germany) with an excitation wavelength of 300–360 nm. Images were analysed and quantified using Fiji software. The percentage of apoptotic cells was calculated using the formula:

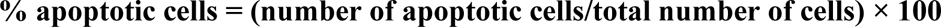

### • ELISA for Protein Markers Detection

To detect the expression levels of protein markers, including TNF-α (Catalogue No. O13850), IL-1β (Catalogue No. O12108), IL-18 (Catalogue No. O12104), CRP (Catalogue No. O11896), CASP-1 (Catalogue No. O10771), IL-6 (Catalogue No. O12137), and tumor suppressor protein (TP53) (Catalogue No. O12769), respective ELISA kits procured from Origin Diagnostics & Research, Kerala, India, were used. ELISA was performed according to the manufacturer’s protocol.

Briefly, protein lysates from various experimental groups of U-87 MG and HepG2 cancer cells were loaded into 96-well plates. After washing away unbound samples with wash buffer, 100 µl of HRP-conjugated secondary antibody was added to each well, followed by a 60-minute incubation at 37°C. Next, 50 µl each of substrate A and substrate B was added to all wells, with a 30-minute incubation at 37°C. The reaction was stopped by adding 50 µl of stop solution to each well, and absorbance was immediately measured at 450 nm using a spectrophotometer (Molecular Devices Spectra-Max M5, USA). Relative protein concentrations were determined by interpolating the absorbance values against a standard curve generated from known concentrations of the standard protein.

### • Immunofluorescence for CD36 Expression Analysis

To perform Immunofluorescence (IF) analysis, U-87 MG and HepG2 cancer cells were initially seeded in 30-mm culture plates at a density of 8×10³ cells per plate. After 24 hours of incubation, the cells were fixed using 4% paraformaldehyde, followed by permeabilization with Triton X-100. Blocking was performed with 3% BSA before incubating the cells with a primary CD36 antibody (Catalogue No. E-AB-33662, Elabsciences, Wuhan, China) at a 1:300 dilution in 3% blocking buffer. After two washes with 1X PBS, the cells were treated with a FITC-conjugated secondary antibody (Catalogue No. E-AB-1014, Elabsciences, Wuhan, China) and counterstained with DAPI. The stained cells were visualized and documented using a fluorescence microscope (Axiovert 40 CFL, Germany).

Additionally, IF was used to evaluate CD36 expression in HepG2 liver cancer cells under treatment with palmitic acid (PA). For this, cells were seeded in 24-well plates and treated with PA at concentrations of 12.5 µg/ml and 50 µg/ml, both individually and in combination with AEV01 at its IC_50_. The experimental protocol was consistent with that used for U-87 MG and HepG2 cells, with fluorescence microscopy employed to capture the stained cells.

Further analysis of CD36 expression was conducted in WRL-68 liver senescence cells using the same IF protocol. Briefly, the cells were seeded, fixed, permeabilized, and blocked, followed by CD36 antibody incubation, secondary antibody treatment, and DAPI counterstaining. Furthermore, evaluation of proliferative ability of these senescence cells with/without AEV01 was executed deploying the similar protocol. Visualization and imaging of the stained cells were performed using the fluorescence microscope (Axiovert 40 CFL, Germany). Following visualization, quantitative analysis was evaluated using ImageJ software.

### • Animal Care and Ethical Considerations

Healthy male Wistar rats (n= 42) weighing between 180 and 240 g, were obtained from Biogen Laboratories, Bangalore, India, for use in this study. The animals were housed in cages under optimal environmental conditions, including a 12:12 hour light-dark cycle, a temperature of 22 ± 1°C, and humidity levels between 50% and 60%. Food and water were provided *ad libitum*. All experimental procedures adhered to the Committee for the Purpose of Control and Supervision of Experiments on Animals (CPCSEA) and Animal Research: Reporting of In Vivo Experiments (ARRIVE) guidelines. The experiments were approved by the Institutional Animal Ethics Committee (IAEC No. 02/IAEC/MGMPRC/01/2023-II, dated 29.8.2023, Sri Balaji Vidyapeeth (Deemed to-be University), Puducherry. Every effort was made to minimize the number of animals used and to reduce their suffering.

### • Stereotactic Implantation of Glioma Cells in Rats

Stereotactic surgery was performed to orthotopically implant glioma cells into healthy rats (n=21), following a standardized protocol ^11,12^. The xenograft recipients were sedated using a ketamine and xylazine cocktail (87 mg/kg and 13 mg/kg body weight, respectively). The skin over the skull was incised from the frontal to the occipital bone to expose the implantation site. The target location for implantation was marked at 3.00 mm lateral and 0.5 mm posterior to the bregma. A small burr hole was drilled at the marked location using a cordless power drill.

Subsequently, 1×10⁶ C6 glioma cells, suspended in 3 µl of PBS, were carefully delivered into the target site at a depth of 6 mm. After the cell injection, the burr hole was sealed with bone wax, and appropriate post-operative care was provided to ensure the well-being of the animals.

### • Subcutaneous Implantation of Hep3B Liver Cancer Cells in Rats

Hep3B liver cancer cells (1×10⁶ cells), resuspended in Dulbecco’s modified Eagle’s medium, were implanted via subcutaneous injection into the right flank of Wistar rats (n=21). The tumors were allowed to develop over a 20-day period until they reached an approximate size of 5 mm in diameter. During this time, the animals were monitored daily for any signs of distress, including weight loss, decreased food and water intake, and dehydration, to ensure their well-being.

### • Grouping and Treatment of Animals Post-Tumor Implantation

Twenty days after tumor implantation, the presence of tumors was confirmed through histopathological examination. Following confirmation, the same procedure was applied to induce tumors in other animals and they were randomly divided into different groups for treatment with the VC and AEV01 as outlined in **Table 1**. The drug dosage was determined based on the previous literature ^13^.

**Table 1:** Animal Grouping and Treatment Plan.

| S.No. | Experimental groups | Glioma<br>(n=21) | HCC<br>(n=21) | Route of<br>administration | Dosage |
| --- | --- | --- | --- | --- | --- |
| 1 | Group I- Healthy control | 6 | 6 | NA | NA |
| 2 | Group II- Tumor Control | 5 | 5 | NA | NA |
| 3 | Group III- Tumor-induced,<br>vehicle-treated | 5 | 5 | Oral | 41.33<br>mg/kg/day |
| 4 | Group IV- Tumor-induced,<br>AEV01-treated | 5 | 5 | Oral | 41.33<br>mg/kg/day |

All the drugs were administered orally for 7 consecutive days. At the end of the treatment, rats were euthanized by asphyxiation using CO_2_, following decapitation. Post euthanasia, brain, liver and other vital organs from the rats were stored in neutral-buffered formalin or −80°C for further analysis.

### • Tissue Preparation and Histological Analysis

After fixation in 10% formalin, brain and liver tissues from all experimental groups were separated, embedded in paraffin, and sectioned to a thickness of 4 µm. The tissue sections were stained with Hematoxylin and Eosin (H&E). Microscopic images of the stained sections

were captured using an inverted microscope (Primo Star, Carl Zeiss, Germany). The images were subsequently analyzed and quantified using ImageJ software.

### • Immunohistochemistry for Assessing Anti-Tumor Effects of AEV01

Immunohistochemistry (IHC) was conducted to evaluate the anti-tumor effects of AEV01 at the molecular level in each experimental group. Tissue sections (5 µm thick) from formalin-fixed samples were prepared. After deparaffinization, antigen retrieval was performed by heating the sections in a microwave for 20 minutes in sodium citrate buffer (10 mM sodium citrate and 0.05% Tween-20, pH 6.0).

To block endogenous peroxidase activity, the tissues were incubated in 3% H₂O₂ for 15 minutes. Subsequently, sections were blocked with 3% bovine serum albumin (BSA) for 10 minutes. Following two washes with 1X-PBST buffer, the slides were incubated with primary antibodies (rabbit anti-CD36, 1:100 dilution) for 1 hour at room temperature. After washing off unbound antibodies, HRP-conjugated secondary antibodies were applied for 30 minutes at room temperature.

Antibody binding was visualized using diaminobenzidine (DAB), and the sections were counterstained with hematoxylin and mounted with mounting medium (Dako, Denmark). Microscopic images were captured using an inverted microscope (Primo Star, Carl Zeiss, Germany). The positive expression of CD36 was assessed and quantified using immunoreactive scores (IRS).

### • ELISA for Protein Marker Detection

ELISA was conducted in the brain and liver tissues samples obtained from the experimental groups of GB and HCC xenograft models, following the same protocol as described in the *in vitro* studies (**Sub-section:** ELISA for Protein Markers Detection).

### • Evaluation of AEV01-Induced Toxicity in Vital Organs

To assess whether AEV01 induced toxicity, the vital organs namely, heart, lungs, liver, kidney, spleen, and pancreas were excised through a mid-line abdominal incision. These organs were then fixed in formalin, embedded in paraffin, sectioned, and subjected to histopathological evaluation. Images of the tissue sections were captured at 10x magnification and analyzed using ImageJ software.

### • Statistical Analysis

All the experiments were conducted in triplicate, and the data were analyzed using GraphPad Prism v.8.0.1 (GraphPad Software, San Diego, CA, USA). Results are presented as Mean ± Standard Deviation (SD). Statistical comparisons between control and experimental groups were performed using Analysis of Variance (ANOVA). Post-hoc for ANOVA was carried out using Dunnett or Tukey’s post-hoc test. p < 0.05 was considered statistically significant.

## Results

### • Cytotoxicity of AEV01 on Cancer and Normal Cell Lines

The cytotoxic effects of AEV01 were evaluated in both U-87 MG GB and HepG2 liver cancer cells using the MTT assay. The drug exhibited a significant, dose-dependent inhibition of cell growth across concentrations ranging from 100 to 700 µg. Notably, the most effective concentrations were observed at 400 µg for GB cells and 300 µg for liver cancer cells, respectively (**Figure 1: A, B**).

**Figure 1:**
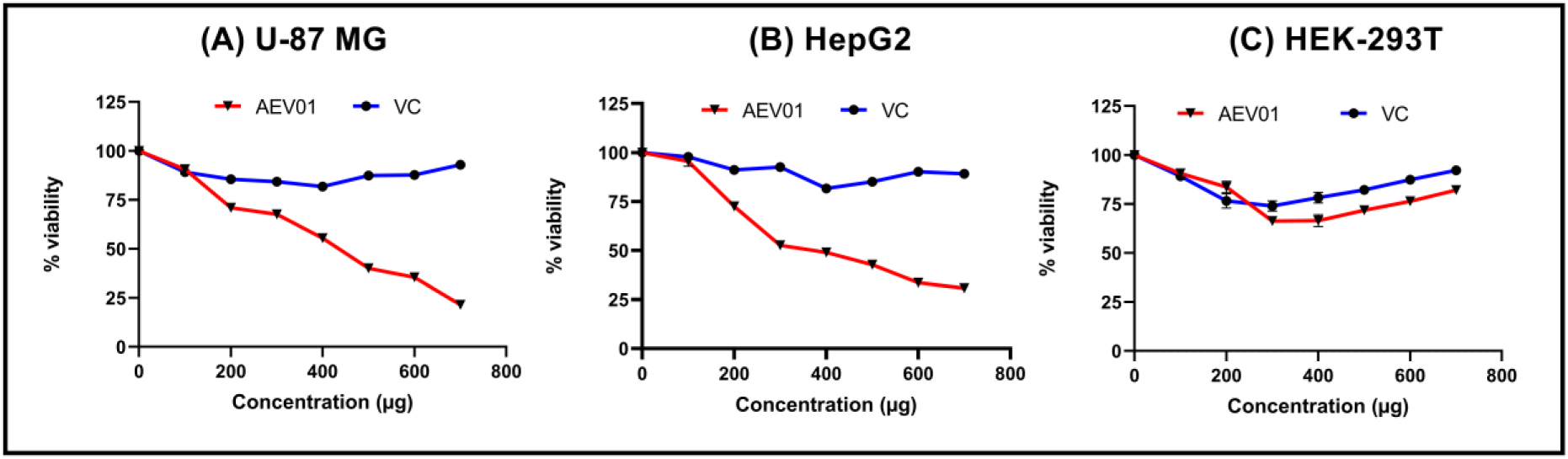
**(A-C)** Cellular viability of malignant GB, liver cancer cells and HEK-293T cells following AEV01 treatment. Cell viability was assessed using MTT assay, n=3.

Further evaluation was conducted on HEK-293T, human embryonic kidney cells, to determine AEV01’s toxicity or tolerance. The MTT assay results (**Figure 1: C**) revealed that AEV01 did not induce significant cytotoxicity in these normal cells. This finding suggests that AEV01 may selectively target cancer cells while exhibiting a tolerable profile in normal non-cancer cells.

### • Induction of Apoptosis by AEV01 in GB and Liver Cancer Cells

Based on the MTT data, AEV01 was found to exert potent growth inhibitory effect in U-87 MG GB and HepG2 liver cancer cells. This conclusion was supported by morphological assessments using AO/EtBr dual staining. In this assay, AO, which permeates live cells, emits green or yellow-green fluorescence in early apoptosis, while EtBr, which stains dead cells, results in orange-red fluorescence indicative of late-stage apoptosis.

From the staining results, it was observed that about 90% of the cells in the control groups were live emitting green fluorescence (**Figure 2: A, B- white arrowheads**). Interestingly, treatment with AEV01 at concentrations of 400 µg and 300 µg significantly induced apoptosis in both U-87 MG and HepG2 cells in comparison with the control (p<0.001) (**Figure 2: A, B- red arrowheads**). The observed apoptosis percentages were 43.2±1.7% for U-87MG cells and 61.4 ±0.8% for HepG2 cells (**Figure 2: A, B**). In contrast, the VC treatment demonstrated that over 90% of the cells remained live in both U-87 MG and HepG2 cultures, indicating that the VC did not significantly induce apoptosis. These morphological findings are consistent with the cytotoxicity data, supporting AEV01’s role as an effective inducer of apoptosis and inhibitor of growth in cancer cells while sparing normal cells.

**Figure 2:**
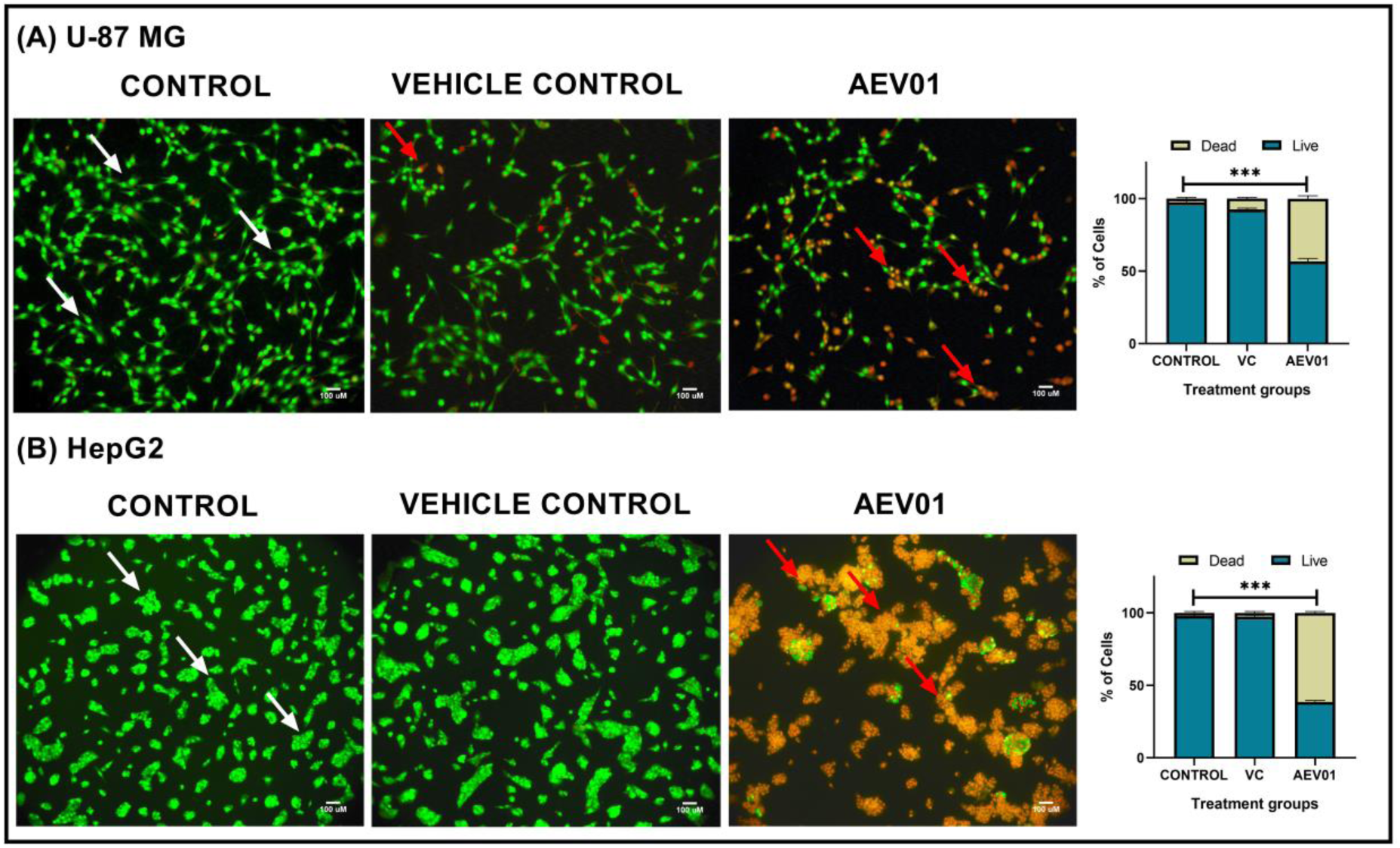
Effect of AEV01 treatment on morphology of GB and liver cancer cells. **(A, B)** U-87 MG and HepG2 cells were treated with AEV01 and stained by AO/EB. Morphological changes in the cells following drug exposure were identified by fluorescent microscopy at 20x magnification (white arrowheads-live cells; red arrowheads-dead cells). Fluorescence intensities were quantified by Fiji software. Data are denoted as Mean ± SD, ***p<0.001 when compared with the control, n=3.

### • Modulation of Inflammatory Markers and Tumor Suppressor Protein by AEV01 in GB and Liver Cancer Cells

In alignment with the significant induction of apoptosis observed in the dual-staining results, the current study further explored the impact of AEV01 at the protein expression level using ELISA. The results demonstrated that AEV01 treatment led to a significant downregulation of the inflammatory markers namely, IL-6, IL-18, IL-1β, TNF-α, CRP, and CASP-1 (p<0.001) in both GB and liver cancer cells (**Figure 3: A-F**). Concurrently, there was an increased expression of TP53 in AEV01-treated cells compared to the control (p<0.001) (**Figure 3: G**). These findings suggest that AEV01 not only inhibits tumor growth and promotes apoptosis but also inhibits cell proliferation, likely through its effects on inflammation and tumor suppression pathways.

**Figure 3:**
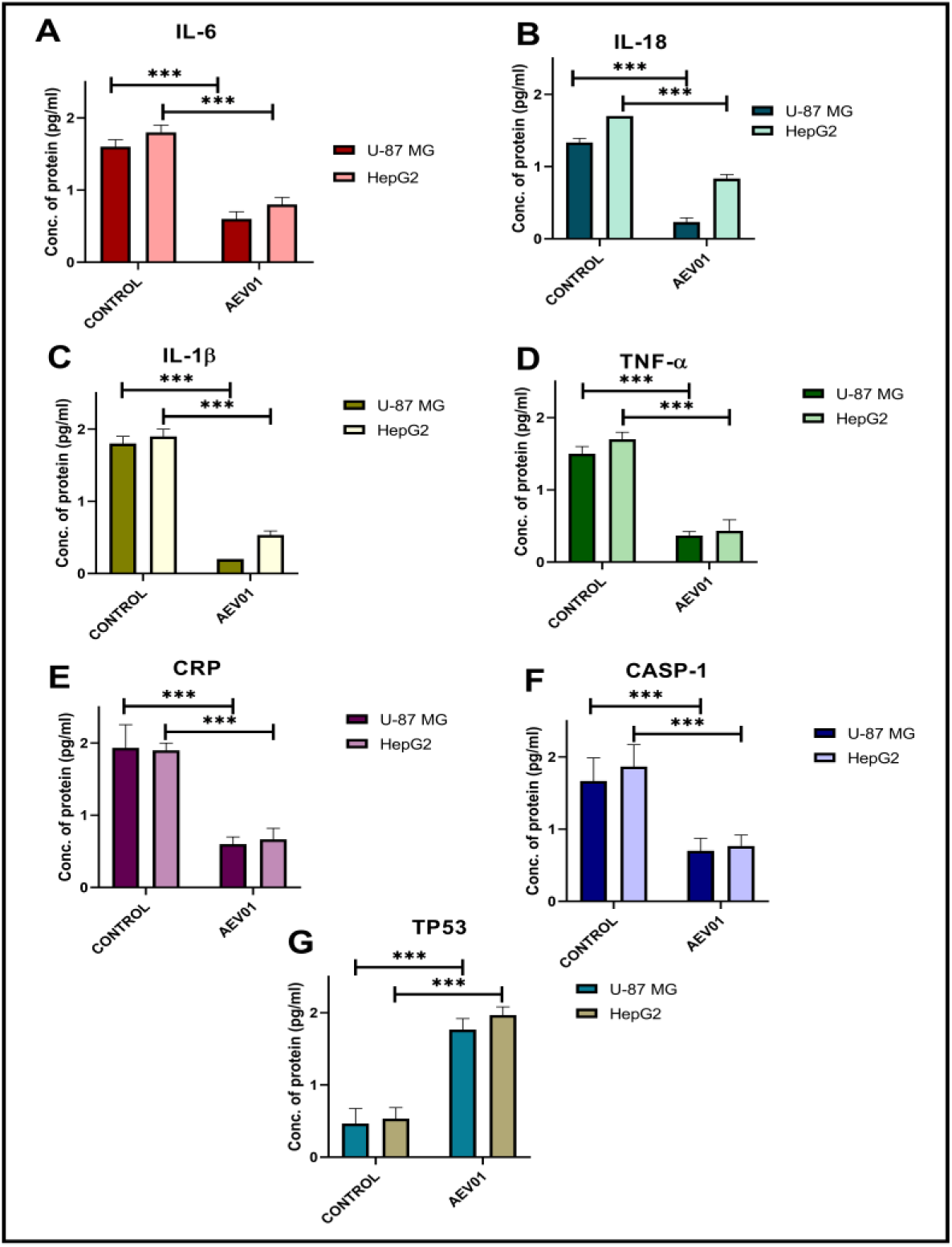
Effect of AEV01 treatment on the protein expression levels of inflammatory and tumor suppressor markers. **(A-F)** Cell lysates of U-87 MG and HepG2 cancer cells treated with AEV01 were analysed for the protein expression levels of IL-6, IL-18, IL-1β, TNF-α, CRP, CASP-1 and **(G)** TP53 by ELISA. Data are denoted as Mean ± SD, ***p<0.001 when compared with the control, n=3.

### • AEV01’s Impact on CD36 Expression Across GB, Liver Cancer, and Senescence Cells

Building upon our preceding observations of AEV01’s impact on inflammatory and tumor suppressor markers, our investigation extended to delve into the potential anti-tumor effects of AEV01, with a specific focus on examining the expression of CD36 in U-87 MG GB and HepG2 liver cancer cells. Employing IF analysis at 20X magnification, CD36 expression was visualized in green, while nuclei were counterstained using DAPI. Notably, a significant downregulation of CD36 expression was observed in both U-87 MG and HepG2 cells following treatment with AEV01, hinting at a potential mechanistic link to the observed anti-tumor effects (**Figure 4: A, B- red arrowheads**).

**Figure 4:**
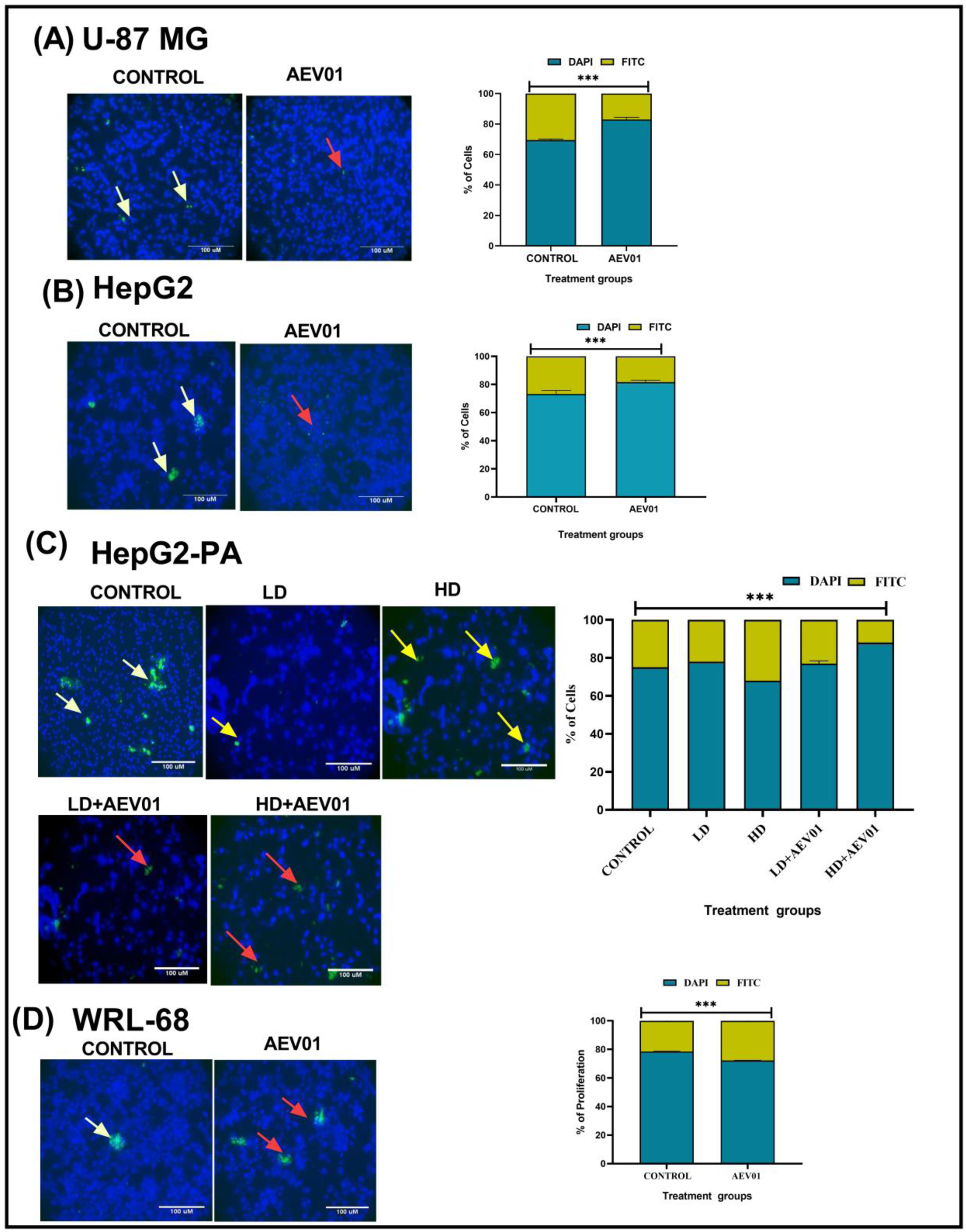
Effects of AEV01 on expression of CD36 **(A)** U-87 MG, **(B)** HepG2, **(C)** Palmitic acid treated HepG2 cells at low and high doses as individually and in combination with AEV01 and **(D)** WRL-68 (senescence cells). Patterns of positive and negative staining for CD36 were identified by fluorescent microscopy at 20x magnification. Quantification was performed using ImageJ software. Data are denoted as Mean ± SD, ***p<0.001 when compared with the control, n=3.

To gain deeper insights into the role of CD36 in HepG2 liver cancer cells, IF analysis was employed to scrutinize the metabolic alterations induced by PA, a critical factor in HCC pathogenesis. Both low and high doses of PA were found to promote CD36 expression (**Figure 4: C-yellow arrowheads).** Intriguingly, the administration of AEV01 (at its IC_50_) in combination with a high dose of PA resulted in a significant downregulation of CD36 compared to the untreated control cells (**Figure 4: C- red arrowheads).**

Furthermore, our study delved into exploring the senescence property induced by CD36 in human liver senescence cells (WRL-68). Utilizing IF analysis at 20X magnification with DAPI counterstaining serving as a negative control, percentage of proliferation was estimated with respect to expression pattern of CD36. In coherence with our earlier findings, AEV01 was found to elevate CD36 expression (35.5%) (**Figure 4: D- red arrowheads)** compared to the control (22%) senescence cells (**Figure 4: D- white arrowheads).** While establishing a direct correlation between CD36 expression and senescence cell growth proved challenging due to limited studies, we interpreted the results in terms of proliferation percentage relative to CD36 expression levels. These intricate findings illuminate a complex interplay between CD36 expression and the effects of AEV01 on both liver cancer cells and senescence cells, prompting the need for further in-depth investigation.

### • Histopathological and CD36 immunostaining effects of AEV01 against GB -induced xenograft model

Extending the thread of our preliminary *in vitro* findings, we sought to substantiate the anti-tumor potential of AEV01 using a glioma-induced animal model. Interestingly, healthy control brain tissues displayed a normal nuclear-cytoplasmic appearance with minimal signs of hemorrhage and very few actively-dividing mitotic cells (**Figure 5: A).** In contrast, the brain tissues from the tumor-control group exhibited a high-index of mitotic cells with intra-haemorrhage lesions and dense nuclei, indicative of aggressive tumor characteristics (**Figure 5: A-black arrowheads).**

**Figure 5:**
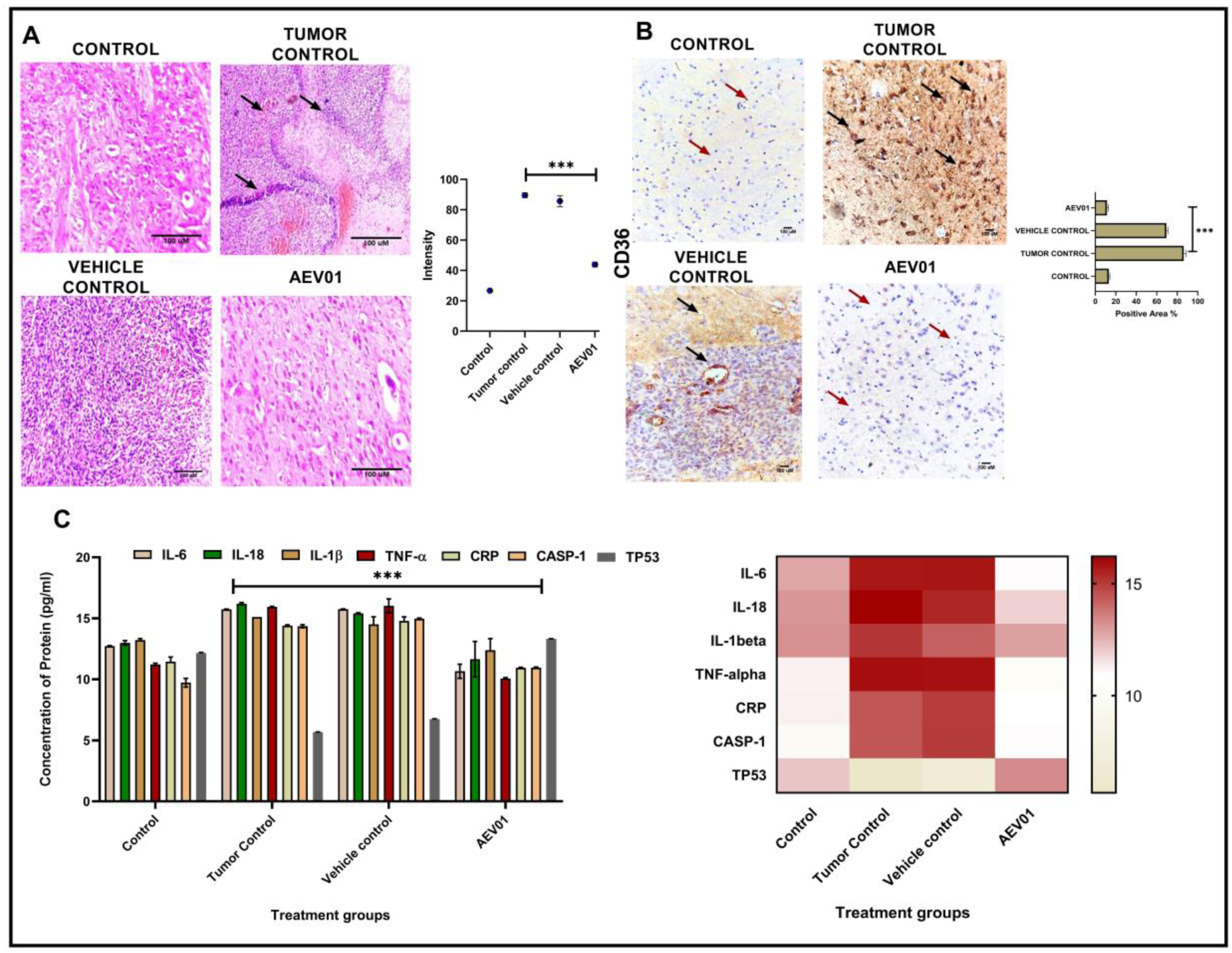
AEV01 treatment-induced histopathological analysis, immunostaining for CD36 and alterations in the protein expression levels of inflammatory and tumor suppressor markers on the brain tissues of glioma xenografts**. (A)** Rat brain tissues treated with AEV01 were subjected to H & E staining. Histopathological changes (black arrowheads) were identified by inverted microscope at 10x magnification. The cellular intensities were quantified using ImageJ software, ***p<0.001 when compared to tumor control. **(B)** Patterns of positive/negative immunostaining for CD36 following administration of AEV01 were identified in inverted light microscope at 40x (positive staining-black arrowheads, negative staining-brown arrowheads). Quantitative data was obtained using ImageJ software, ***p<0.001 when compared to the tumor control, n=3. (C) Brain tissues of the GB-bearing xenografts, following administration AEV01 were analysed for the protein expression levels of inflammatory and tumor suppressor markers by ELISA. Data are represented as Mean ± SD, ***p<0.001 when compared to tumor control, n=3.

Interestingly, AEV01-treated glioma-induced xenograft tissues demonstrated a morphological appearance similar to that of healthy control tissues (p<0.001). This contrasted with the tumor-control group, showcasing a lower mitotic index and an absence of hemorrhage signs (**Figure 5: A).** Notably, the vehicle control did not exert any significant effect on tumor growth (**Figure 5: A).** These findings strongly suggest that AEV01 has the potential to restore the morphological appearances of tumor xenografts with minimal side effects.

In alignment with these *in vivo* observations, we further explored the immunohistochemical expression of CD36-a fatty acid translocase protein with implications in lipid metabolism and cellular reprogramming in cancer cells, including glioma. The IHC analysis of CD36 was conducted across each experimental group. In healthy control brain tissues, CD36 expression was minimal (**Figure 5: B-brown arrowheads)**, while the tumor-control group exhibited elevated CD36 expression, evident by cytoplasmic positivity (**Figure 5: B-black arrowheads).** Interestingly, AEV01-treated glioma tissues significantly downregulated the expression of CD36 compared to both tumor-control and vehicle-control experimental groups (p<0.001) (**Figure 5: B-brown arrowheads).**

The cytoplasmic positivity of CD36 observed in AEV01-treated glioma tissues mirrored that of healthy control-treated groups (**Table 2**). Overall, AEV01 demonstrated a reduction in tumor growth formation, possibly mediated through the downregulated expression of CD36, in comparison to the tumor-control group, as supported by the IHC analysis. These results reinforce the potential therapeutic efficacy of AEV01 in glioma treatment, validated through both *in vitro* and *in vivo* studies.

**Table 2:** Immunohistochemical scores of CD36 against each of the experimental groups.

| Treatment groups | Intensity | % Positivity | IHC scores<br>(intensity × %<br>positivity) |
| --- | --- | --- | --- |
| Healthy control | 0 | 1 | 0 |
| Tumor-control | 2 | 2 | 4 |
| AEV01-treated | 1 | 2 | 2 |
| Vehicle control | 2 | 2 | 4 |

### • AEV01’s Inhibitory Effects on GB Progression Through Modulation of Inflammatory Markers and TP53 Expression

In alignment with the compelling *in vivo* pathogenesis study outcomes, where AEV01 showcased a reduction in tumor growth formation and CD36 expression, we extended our investigation to the protein expression level using ELISA. The examination revealed that AEV01-treated glioma tissues demonstrated a downregulation in the expression of inflammatory markers, including IL-6, IL-18, IL-1β, TNF-α, CRP, and CASP-1 (p<0.001), alongside an elevated expression of TP53, in comparison to healthy control tissues (**Figure 5: C**). These results indicate that AEV01 hinders the propagation of tumor growth through the downregulation of inflammatory markers associated with inflammasome formation. Simultaneously, there is a concurrent upregulation of TP53, a crucial tumor suppressor marker. The consistency across multiple levels of analysis, from histopathology to IHC and now to ELISA, further strengthens the evidence supporting AEV01’s inhibitory effects on glioma progression.

### • Histopathological effects of AEV01 against HCC-induced animal model

Besides elucidation of the anti-glioma potency of AEV01 *in vivo*, the present study aimed to explore the anti-hepatocellular potential of AEV01 in an Hep3B-induced HCC model. As per our preliminary *in vitro* findings, AEV01 significantly altered the morphology of liver cancer cells and induced apoptosis. As a step towards further validation, the anti-cancer potential of AEV01 was assessed in Hep3B-induced HCC xenografts. As per the histopathological findings, tumor control-liver tissue expressed formation of hemorrhagic lesions and poorly differentiated hepatocyte appearance, whereas, AEV01 treated tissues reverted towards a normal hepatocyte appearance (**Figure 6: A-black arrowheads),** while vehicle control did not exert any alterations in the tumor architecture and hence its efficacy is undermined to that of treated tissue (**Figure 6: A).** Interestingly, upon exposure to AEV01 in Hep3B-induced HCC model, the tissue architecture was restored to that of normal healthy control as shown in **Figure 6: A**.

**Figure 6:**
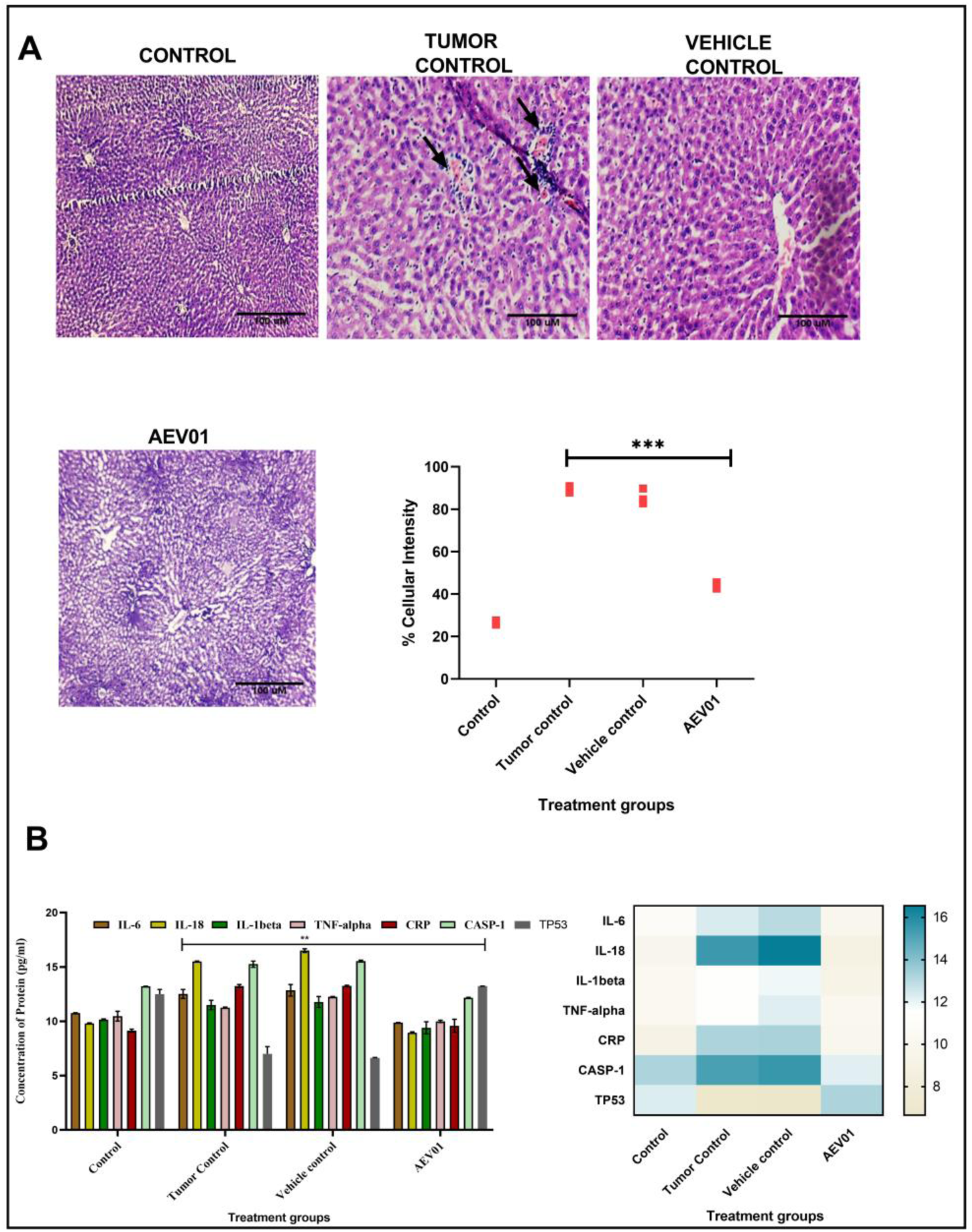
AEV01 treatment-induced histopathological analysis, and alterations in the protein expression levels of inflammatory and tumor protein markers on the liver tissues of HCC xenografts**. (A)** Rat liver tissues treated with AEV01 were subjected to H & E staining. Histopathological changes (black arrowheads) were identified by inverted microscope at 10x magnification. The cellular intensities were quantified using ImageJ software, ***p<0.001 when compared to tumor control. **(B)** Liver tissues of Hep3B-induced HCC xenografts, following administration AEV01 were analysed for the protein expression levels of inflammatory and tumor suppressor markers by ELISA. Data are represented as Mean ± SD, ***p<0.001 when compared to tumor control, n=3.

### • Validation of *in vitro* findings through ELISA analysis in HCC-induced animal models

Following, exposure of HCC-induced tumor animals to AEV01 treatment, proteins isolated from tissue samples was subjected to ELISA in order to validate the *in vitro* findings. Similar to our *in vitro* findings, it was observed that expression levels of the inflammatory markers namely, IL6, IL-18, IL-1β, TNF-α, CRP and CASP-1 was reduced in AEV01-treated rats, whereas TP53 expression was elevated when compared to tumor and VC-treated rats significantly (**Figure 6: B**).

### • Histopathological Analysis Shows AEV01’s Safety and Anti-Tumor Effects against GB and liver cancers

The preliminary study findings focused on assessment of anti-tumor potential of AEV01 against GB and HCC cancer cells, wherein, apoptosis was triggered in these cancer cells mediated via downregulated expression of CD36, a key component contributing towards tumorigenesis process. On grounds of these, *in vivo* experiments were conducted, whereby, we observed that AEV01 impacted the growth of tumor mediated via downregulation of inflammatory markers against GB and HCC-induced xenograft models. We observed that AEV01 significantly impacted tumor growth against both of these models.

However, anti-tumor drugs/treatment are assumed to exert adverse side effects, thereby, histopathological analysis was performed to assess if any toxicity was induced against the vital organs. Interestingly, histopathological analysis of the heart and lungs from all the glioma-induced experimental groups depicted normal cardiac muscle fibres and alveolar structures without any edema lesions (**Figure 7**). Similarly, liver tissues (glioma model) portrayed clear hepatocytes structures without any signs of fatty deposits or necrosis (**Figure 7**). Histopathological sections of kidney depicted clear and distinct visible nucleolus, without any necrotic infiltrations against all of the experimental groups (**Figure 7**). Also, photomicrographs of spleen and pancreas did not exhibit any abrupt alterations in their morphological appearance (**Figure 7**).

**Figure 7:**
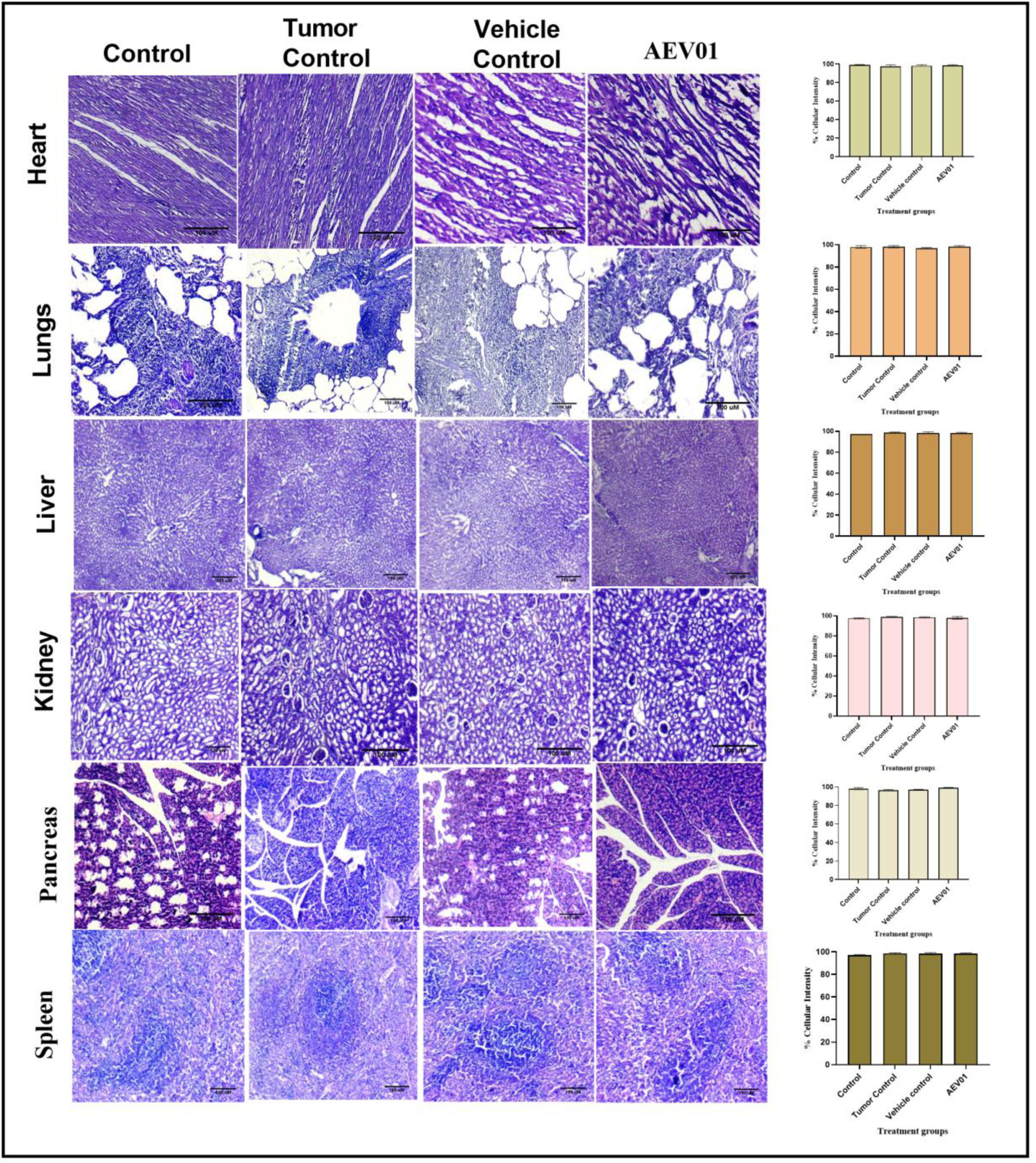
Histopathological changes following administration of AEV01 on the vital organs of glioma-induced rats such as heart, lungs, liver, kidney, spleen, and pancreas were identified by inverted microscope at 10x magnification. Quantitative data based on the cellularity index was obtained employing ImageJ. Data are represented in terms of Mean ± SD, n=3.

Likewise, histopathological observation was performed in the vital organs obtained from all the experimental groups of HCC-induced animal model (**Figure 8**). Interestingly, all the vital organs namely, heart, lungs, pancreas, spleen and kidney did not exhibit any abrupt alterations in their morphological appearance (**Figure 8**). Taken together, our study indicates that AEV01 is tolerable at its respective doses, while inducing pathological effect against both GB and HCC-induced xenograft models (**Figure 7 & 8).**

**Figure 8:**
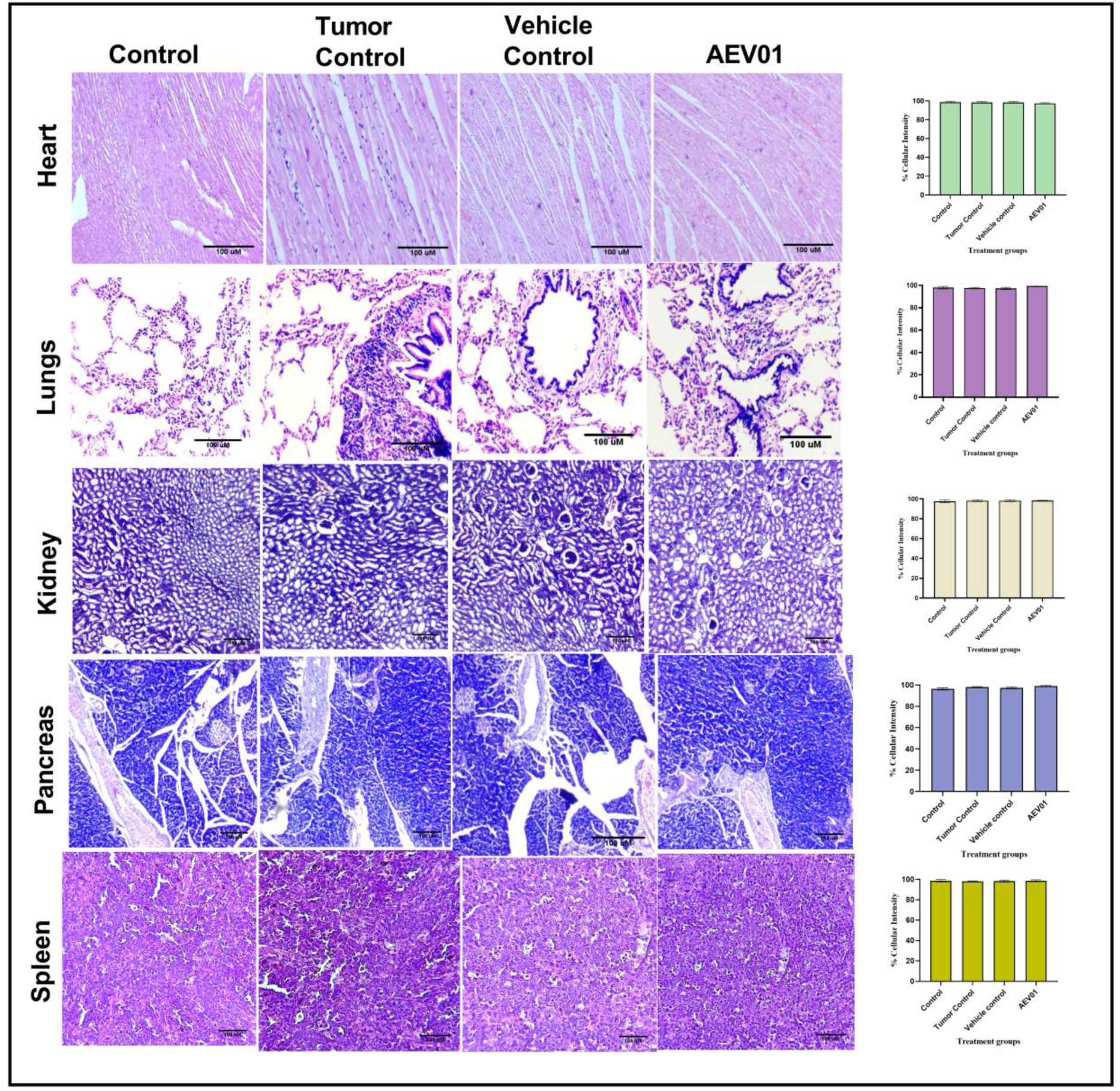
Histopathological changes following administration of AEV01 on the vital organs of HCC-induced rats such as heart, lungs, kidney, pancreas and spleen were identified by inverted microscope at 10x magnification. Quantitative data based on the cellularity index was obtained employing ImageJ. Data are represented in terms of Mean ± SD, n=3.

## Discussion

Cancer is considered as one of the leading causes of death worldwide, due to its complex biology and the limitations of current therapies ^14^. The rising concern of mortality cases due to cancer has necessitated the dire need to implement alternative therapeutic strategies ^15^. The results of the present study highlight the potential of AEV01 as a promising therapeutic agent for GB and HCC.

AEV01, derived from the extract of *Picrorhiza kurroa*, has demonstrated significant cytotoxic effects against both U-87 MG GB and HepG2 liver cancer cells. This was evidenced by the dose-dependent inhibition of cell growth with effective concentrations being 400 µg and 300 µg, respectively. Interestingly, the MTT assay results revealed that AEV01 did not induce significant cytotoxic effects in HEK-293T cells, suggesting that AEV01 might preferentially target cancer cells over normal cells (**Figure 1: A-C**). This finding aligns with the therapeutic aim of minimizing off-target effects and preserving normal tissue function while combating tumor growth. Such findings corroborate with previous research^16,17^.

Further, morphological examination of apoptosis induction by AEV01 on U-87 MG and HepG2 cells was conducted using AO/EtBr dual stain. AO, being permeable to live cells, appears green, while cells in the early stage of apoptosis exhibits yellow-green fluorescence. On the other hand, EtBr stains dead cells, particularly those in the late stage of apoptosis, producing an orange-red fluorescence ^18^. In this study, the treatment with AEV01 on both U-87 MG and HepG2 cells significantly induced apoptosis, with percentages measured at 43.2±1.7% and 61.4 ±0.8% respectively (**Figure 2: A, B)**. Remarkably, the VC (**Figure 2: A, B)** demonstrated that over 90% of cells in both U-87MG and HepG2 cells remained viable. This observation indicates that the VC did not elicit apoptosis in these cells. The lack of apoptosis induction in the vehicle control provides a crucial baseline for comparison with the AEV01-treated groups. This result suggests that any observed apoptotic effects in the AEV01-treated cells are specific to the treatment and not merely a consequence of the experimental conditions or vehicle used. The absence of significant apoptosis in the VC groups underscores the specificity of AEV01 in triggering apoptotic pathways in U-87 MG and HepG2 cells. These findings enhance the credibility of the observed apoptotic effects in the treated groups, reinforcing the idea that AEV01 possesses anti-tumor properties by inducing programmed cell death in these cancer cells.

In line with the previously observed significant induction of apoptosis, the current study further explored the modulation of inflammatory and tumor suppressor markers, revealing a comprehensive mechanism by which AEV01 exerts its anti-cancer effects. Specifically, the downregulation of inflammatory markers such as IL-6, IL-18, IL-1β, TNF-α, CRP, and CASP-1 suggests that AEV01 effectively disrupts the pro-inflammatory environment that often supports tumor growth and survival (**Figure 3: A-F).** This reduction in inflammation likely contributes to the compound’s ability to halt tumor progression. Additionally, the upregulation of TP53, a crucial tumor protein marker, in AEV01-treated cells points to the activation of important pathways that further inhibit tumor cell proliferation (**Figure 3: G).** The increased expression of TP53 not only complements the apoptotic effects but also strengthens the overall anti-cancer strategy by promoting cell cycle regulation and preventing uncontrolled cell growth. This dual action-suppressing inflammation and enhancing tumor suppression-demonstrates AEV01’s potential as a powerful therapeutic agent in targeting both the survival and proliferation mechanisms of glioma and liver cancer cells.

Based on the previous findings that highlighted AEV01’s ability to modulate inflammatory and tumor suppressor markers, its impact on CD36 expression further elucidates its potential anti-tumor mechanisms. CD36, a fatty acid translocase, is strongly associated with the progression of HCC by promoting tumor growth, proliferation, migration, and invasion ^19^. In addition, CD36 also alters the metabolic state of cancer cells by favoring oncogenic glycolysis over oxidative phosphorylation and is linked to a tumor-promoting phenotype characterized by immunosuppressive and pro-inflammatory activities ^20^. Elevated levels of CD36 are correlated with increased tumor-infiltrating macrophages, that suppress anti-inflammatory responses, making CD36 a promising therapeutic target in cancer treatment ^20^. In this study, IF analysis revealed a significant downregulation of CD36 in U-87 MG GB and HepG2 liver cancer cells after AEV01 treatment, suggesting a plausible mechanism for its anti-tumor effects (**Figure 4: A, B).** Moreover, further investigation in liver cancer cells demonstrated that AEV01, when combined with PA, a factor crucial in HCC pathogenesis, significantly reduced CD36 expression (**Figure 4: C).** This reduction might be associated with the lipotoxicity induced by PA, leading to pyroptosis, as evidenced by the upregulated expression of caspase-1 and IL-1β ^21^. While apoptosis through this mechanism is evident, further validation at the transcriptional and translational levels is needed to fully understand the pathways involved.

Additionally, the study explored CD36 expression in human liver senescence cells (WRL-68) using IF analysis. AEV01 was observed to elevate CD36 expression (35.5 %) compared to the control (22%), possibly influencing the proliferation capacity of senescence cells (**Figure 4: D).** Although a direct correlation between CD36 expression and senescence cell growth could not be established, the findings suggest that increased expression of CD36 in senescence cells, exposed to treatment could might have a complex role in tumorigenesis. This complexity arises from the dual nature of senescence, which can either downregulate tumor suppressor genes or provoke the synthesis of proto-oncogenes^22^. Thus, the downregulation of CD36 observed in GB and liver cancer cells, particularly in the context of metabolic alterations induced by the AEV01 and PA combination, presents a promising avenue for anti-tumor therapeutic interventions.

Building on our initial *in vitro* findings, which showed AEV01’s potential anti-tumor effects by suppressing inflammatory markers and promoting apoptosis, we validated these results using an orthotopic xenograft-induced glioma and HCC model. Previous studies have shown that *Picrorrhiza kurroa* extracts was able to suppress tumor formation and extend lifespan in various tumor-induced animal models, supporting their potential as anti-tumor agents ^17^. In alignment with these findings, AEV01-treated GB tissues, in this study, exhibited a morphological appearance similar to that of healthy controls, with a lower mitotic index and an absence of haemorrhage (**Figure 5: A).** This was in contrast to the tumor control group, which showed dense nuclei, intra-haemorrhage lesions, and a high mitotic cell count. Furthermore, immunohistochemical analysis revealed that AEV01 downregulated CD36 expression, a protein associated with tumorigenesis and lipid metabolism, when compared to both the tumor and vehicle-control groups (**Figure 5: B, Table 1).** This downregulation was significant and correlated with reduced tumor growth, suggesting that AEV01 not only inhibits tumor progression but also potentially restores the tissue’s normal morphology with minimal side effects. ELISA analysis of tissue samples revealed a reduction in inflammatory markers expression (IL-6, IL-18, IL-1β, TNF-α, CRP, and CASP-1) and an elevation in TP53 expression following AEV01 treatment (**Figure 5: C).**

In consensus with the previous findings, the Hep3B-induced tumor control exhibited hemorrhagic lesions and undulating tumor architecture, whereas, AEV01 treated tissue sample restored the morphological appearance of hepatic tissues was restored to that of control (**Figure 6: A**). Moreover, ELISA demonstrated a downregulation of the inflammatory markers, IL-6, IL-18, IL-1β, TNF-α, CRP, and CASP-1, with concomitant upregulation of TP53 following AEV01 exposure (**Figure 6: B).** The downregulation of inflammatory markers indicates that AEV01 may interfere with the chronic inflammation often associated with tumor progression, thereby disrupting the tumorigenic environment. In parallel, the upregulation of TP53, a crucial regulator of cell cycle and apoptosis, underscores AEV01’s role in promoting tumor cell death and preventing malignant proliferation. These findings are particularly significant in the context of GB and HCC, cancer types known for their complex etiology involving chronic inflammation and disrupted apoptotic pathways. Thus, the ability of AEV01 to address both of these aspects simultaneously suggests that it could offer a more effective and targeted approach compared to current therapies, which often focus on a single pathway or mechanism.

Moreover, this study also showed that AEV01 has a good safety profile. There were no significant abnormalities observed in important organs such as the heart, lungs, liver, kidneys, spleen, and pancreas in both GB and HCC xenografts (**Figure 7 & 8).** This lack of harmful side effects, along with its targeted action against cancer cells, suggests that AEV01 might be a safer option compared to traditional chemotherapy or radiation therapy, which often has severe side effects because it affects both cancerous and healthy cells.

Although our *in vitro* and *in vivo* studies, along with unpublished data from pancreatic cancer and COVID-19 human clinical studies, have demonstrated AEV01’s safety and efficacy, further research is essential. Future studies should focus on elucidating the precise molecular mechanisms underlying AEV01’s anti-tumor effects and evaluating its long-term safety and efficacy in preclinical and clinical settings. Comprehensive pharmacokinetic and pharmacodynamic studies are also essential to optimize dosing regimens and maximize therapeutic benefits while minimizing potential adverse effects. Overall, these findings contribute to the growing evidence supporting AEV01’s therapeutic potential in cancer treatment and highlight the need for continued research.

In conclusion, this study emphasizes the promising potential of AEV01 as a therapeutic agent for GB and HCC. Derived from *Picrorhiza kurroa*, AEV01 demonstrated significant cytotoxic effects against cancer cells, effectively targeting both U-87 MG GB and HepG2/Hep-3B liver cancer cells while showing minimal toxicity to normal cells. The induction of apoptosis in these cancer cells, coupled with the reduction in inflammatory markers and upregulation of TP53, highlights AEV01’s dual mechanism of action by suppressing inflammation and enhancing tumor suppression. Additionally, the downregulation of CD36, a protein associated with tumor progression and metabolic alterations, further supports AEV01’s potential as an anti-cancer agent. The favorable safety profile observed in vital organs, alongside its specific targeting of cancer cells, suggests that AEV01 may offer a safer alternative to conventional therapies with fewer off-target effects (**Figure 9**).

**Figure 9:**
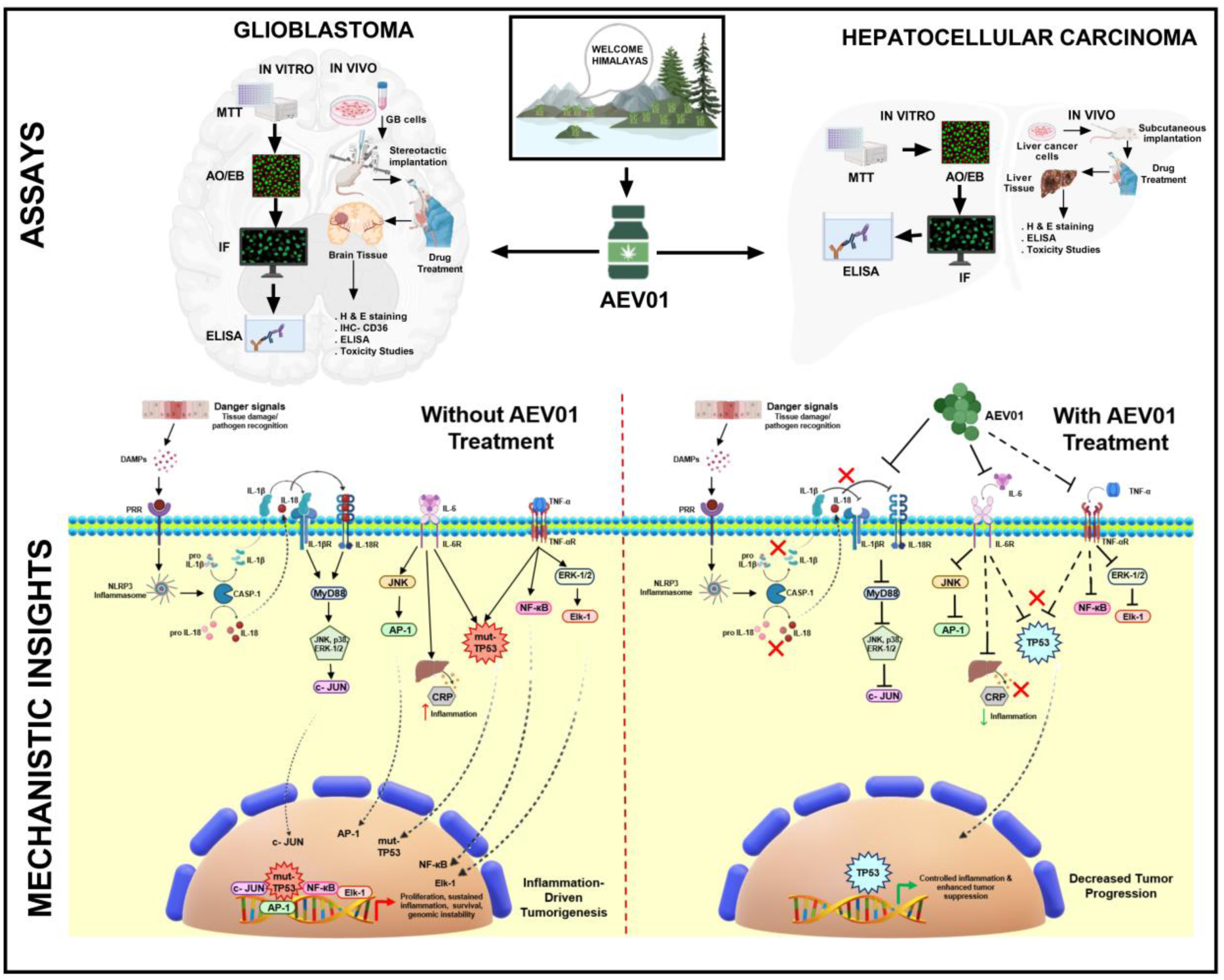
Schematic representation of the workflow and mechanism of action of AEV01 on glioma and HCC models *in vitro* and *in vivo*.

Overall, the comprehensive evidence from *in vitro* and *in vivo* studies positions AEV01 as a promising candidate for further clinical development in cancer therapy. Its ability to address both inflammation and tumor growth mechanisms, while maintaining a good safety profile, warrants continued research into its therapeutic potential and mechanisms of action.

## Acknowledgements

The authors thank the Department of Biochemistry and Molecular Biology, Pondicherry University for accepting and carrying forward this research work as a Consultancy Project and Dr. Agieshkumar B, Mahatma Gandhi Medical Advanced Research Institute (MGMARI), SBV for supporting the animal studies associated with the consultancy.

## Author Contributions Statement

ATS and MMS conceived the experiments and reviewed the manuscript text, IB & D.P.S conducted the experiments and wrote the manuscript, MMS funded the research work to understand the molecular mechanism, AK and RM analysed the results. All authors reviewed the manuscript.

## Competing Interests

The authors declare that M/s. Astrel Genome Ltd, India is involved in the current research work, due to their knowledge on the novel product, AEV01.

## Notes

### Competing Interest Statement

The authors have declared no competing interest.

### Summary of Updates

The abstract and the Introduction part were updated regarding the effect of AEV01 in glioma and hepatocellular carcinoma cells.

